# Differential chromatin response to retinoic acid in neuroblastoma according to type of *ATRX* mutation

**DOI:** 10.1101/2025.07.14.664731

**Authors:** Federica Lorenzi, Matthew Shipley, Luke Deane, Robert Goldstone, Vidur Tandon, Barbara Martins da Costa, Kevin Greenslade, Karen Barker, Fariba Nemati, Angela Bellini, Gudrun Schleiermacher, Louis Chesler, Francois Guillemot, Sally L George

## Abstract

Neuroblastoma is a childhood cancer, arising in the developing sympathetic nervous system. Differentiation therapy with 13-cis-retinoic acid (RA) is routinely given to children with high-risk neuroblastoma in the minimal residual disease setting to prevent relapse, however there is little understanding of which patients benefit from RA therapy.

*ATRX* alterations are identified in 10% of high-risk neuroblastomas and associated with poor outcomes. The commonest type of *ATRX* alterations in neuroblastoma are in-frame multi-exon deletions, followed by nonsense mutations predicted to result in loss-of-function (*ATRX* LoF).

We treated paired *ATRX* wild-type and *ATRX* LoF neuroblastoma cell lines with RA and show that cells with *ATRX* LoF fail to upregulate direct RA target genes. Cells with *ATRX* LoF also show reduced chromatin accessibility at genes involved in differentiation and development following RA treatment. Conversely, neuroblastoma models with in-frame deletions mount a response to RA and show *in-vitro* sensitivity to RA. Taken together this shows that the mechanism of differentiation in *ATRX*-altered neuroblastoma depends on the type of *ATRX* alteration, with implications relating to both oncogenesis and therapeutic response.

## INTRODUCTION

Neuroblastoma is a heterogenous cancer of the sympathetic nervous system, arising in cells of neural crest lineage during embryonal development. It can spontaneously differentiate and regress, however more frequently it presents in young children as an aggressive, widely metastatic malignancy. A limited number of genetic alterations have been identified as drivers of neuroblastoma, including somatic *ATRX* gene alterations which are identified in 10% of high-risk neuroblastomas and define a distinct sub-group of patients with chronic, slowly progressive disease [1-3].

*ATRX* codes for a chromatin remodeling protein with far reaching functions. It has multiple roles including the maintenance of genomic stability at repetitive DNA regions and the resolution of R-loops, G quadruplexes and stalled replication forks. ATRX is also known to regulate cell-state specific gene expression via multiple different mechanisms [4].

The most frequent type of *ATRX* alterations seen in neuroblastoma are in-frame multi-exon deletions, most commonly resulting in the loss of exons 2-10 of the gene (containing the EZH2 and ADD domains), and resulting in an in-frame fusion (IFF) protein [5, 6]. The remainder of *ATRX* alterations are either nonsense mutations predicted to result in loss of function (LoF) (18%), and missense mutations, which tend to cluster in the helicase domain of the gene (14%) [7, 8]. These different types of alterations are all universally associated with certain biological characteristics such as the presence of alternative lengthening of telomeres [5] and an immunogenic phenotype [9]. However, evidence is mounting that there are also significant differences between *ATRX* IFF and *ATRX* LoF neuroblastoma in the underlying mechanisms of epigenetic deregulation [6, 7]. In *ATRX* IFF neuroblastoma it has been shown that differentiation block is mediated by a non-canonical function of the *ATRX* IFF, activating REST and resulting in repression of neurogenesis genes [6]. However, how *ATRX* LoF affects neuroblastoma differentiation potential is unknown.

Here, we show impairment of the response to differentiation stimuli in both stem cell and neuroblastoma models with ATRX LoF. Neuroblastoma cells with ATRX LoF also show decreased chromatin accessibility at neuronal differentiation genes following treatment with 13-*cis*-retinoic acid (RA) - an agent used as standard-of-care in the minimal residual disease setting to prevent neuroblastoma relapse. Conversely, *ATRX* IFF neuroblastoma models are broadly sensitive to RA and show appropriate opening of chromatin at RA response sites. Taken together this work shows that the mechanism of differentiation block in neuroblastoma depends on the type of *ATRX* alteration, with implications relating to both oncogenesis and therapeutic response.

## RESULTS

### *ATRX* LoF neuroblastoma cells have an impaired chromatin response to RA compared to their wild-type counterparts

To investigate the role of *ATRX* in differentiation, we used paired *ATRX* wild-type (*ATRX* WT) and LoF induced pluripotent stem cells (iPSCs) [10] to generate human axial progenitor (HAP) cells, as the first step in differentiation towards neural crest lineage (**Fig S1A-E**) [11]. Upon HAP differentiation, we saw a robust induction of expression of the axial progenitor marker *CDX2* in both *ATRX* WT and LoF iPSCs (**Fig S1F**), however this was only associated with induction of *HOXC* gene expression in *ATRX* WT cells (**Fig S1G**).

In normal embryonic development, CDX2 co-ordinates the spatial and temporal expression of *HOX* genes together with RA [12, 13]. Given that RA is also used clinically as a differentiation therapy for patients with neuroblastoma, we next asked if a similar phenotype was seen when *ATRX* LoF neuroblastoma cells were treated with RA. Paired *ATRX* WT (p53(2)) and *ATRX* LoF (E6) neuroblastoma cells [14] were treated with RA and evaluated for expression of known RA target genes: *CYP26A, HOXA1* and *HOXA4* [15]. Concordant with our findings in *ATRX* LoF iPSCs, we also identified a failure of upregulation of direct RA target genes in neuroblastoma cells with *ATRX* LoF (**Fig 1A-B**).

**Figure 1.**
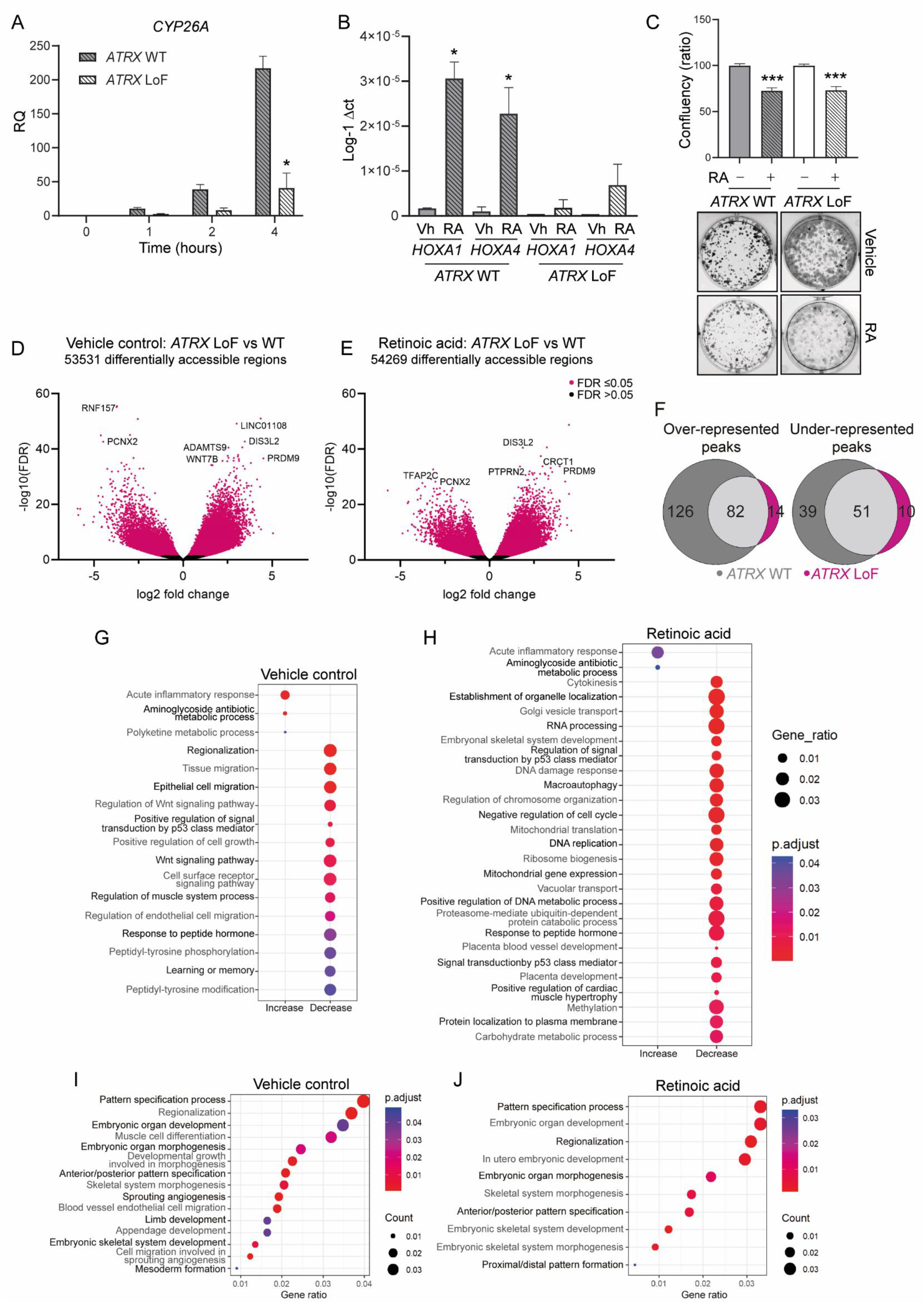
*ATRX* LoF neuroblastoma cells have an impaired chromatin response to RA compared to their wild-type counterparts. **(A)** Time course experiment showing expression of *CYP26A*, normalized to *GAPDH* following exposure of p53(2) (*ATRX* wildtype) and E6 (*ATRX* LoF) cells to retinoic acid (RA). **(B)** *HOXA1* and *HOXA4* expression by RT-qPCR following 72 hours treatment with RA, compared to vehicle control. Inverse log delta CT is indicated. Where no expression was seen, this is indicated as 0. For all RT-qPCR experiments: *t* test, \**P* < 0.05. **(C)** Clonogenic assay results for p53(2) and E6 cell lines following treatment with 10µM RA versus control (*t* test, \*\*\**P* < 0.001). **(D-E)** Volcano plots showing differentially accessible regions in E6 versus p53(2) cell lines following **(D)** vehicle control treatment conditions and **(E)** RA treatment. **(F)** Venn diagrams comparing differentially accessible transcription factor binding motifs following RA treatment in p53(2) and E6 cell lines. **(G-J)** Gene ontology analysis, comparing differential accessibility at gene promoters between E6 and p53(2) lines in **(G)** vehicle control treated conditions and **(H)** following treatment with RA. Displayed terms have been selected using REVIGO which eliminates redundant GO terms. **(I-J)** List of pathways grouped under the term “regionalization” and “embryonic skeletal system development” by Revigo for **(I)** vehicle control and **(J)** RA treated samples, respectively.

We next analysed sensitivity to RA in the paired cell lines by clonogenic assay. We saw no difference in sensitivity to RA between *ATRX* WT and LoF cells (**Fig 1C**) however, the parental cell line was already highly retinoic acid resistant. Of note, the parental cell line is derived from a SH-EP cell [14] which is known to be a mesenchymal cell line with a super-enhancer landscape that associated with RA resistance [16]. Therefore, we decided to evaluate the direct effects of ATRX LoF on chromatin accessibility by ATAC sequencing **(Fig 1D-J)**.

We identified widespread differences in chromatin accessibility in *ATRX* LoF compared with *ATRX* WT neuroblastoma in both vehicle control conditions (**Fig 1D, Table S1**) and upon treatment with RA **(Fig 1E, Table S1)**. Transcription factor binding motif analysis of differentially accessible peaks in response to RA also identified that a greater number of both over and under-represented peaks were seen in response to RA in *ATRX* WT cells compared with *ATRX* LoF cells (**Fig 1F, Table S1**).

We performed gene ontology (GO) analysis of differential accessibility in gene promoter regions between ATRX LoF and WT cells. Given the large number of terms identified, we used REVIGO [17] to summarize and remove redundant GO terms (**Fig 1G-H**). The key GO terms with an increase in accessibility in ATRX LoF cells were in acute inflammatory response pathways (**Fig 1G-H**), consistent with our work identifying an immunogenic phenotype in *ATRX-*altered neuroblastoma [9]. Many GO terms with decreased accessibility in *ATRX* LoF cells were identified, both in vehicle control and RA treated conditions, relating to multiple cellular processes (**Fig G-H**). We also evaluated for differences in individual GO terms related to differentiation between *ATRX* LoF and WT cells which identified decreased accessibility in multiple critical pathways in embryonic development in *ATRX* LoF cells compared with WT cells in both vehicle control and RA treated conditions (**Fig 1I-J, Table S1**).

### Maintenance of epigenetic response to RA is seen in neuroblastoma cell lines with *ATRX* IFF’s

We next assessed whether RA treatment induced expression of direct RA *HOX*-gene targets in two neuroblastoma cell lines with in-frame *ATRX* deletions: SK-N-MM and CHLA-90 [6]. In contrast to *ATRX* LoF cells, cell lines with the *ATRX* IFF were able to upregulate direct RA *HOX*-gene targets following treatment with RA (**Fig 2A**). We then performed ATAC sequencing in the *ATRX* IFF cell lines to evaluate changes in chromatin accessibility following RA treatment **(Fig 2B)**. Large changes in chromatin accessibility in response to RA were seen in SK-N-MM (**Fig 2C, Table S2**) and more limited changes in chromatin accessibility were seen in CHLA-90 (**Fig 2D, Table S2**). Although comparisons could not be made with matched wild-type cells in these models, we noted that both cell lines showed highly significant increases in chromatin accessibility at the direct RA genes *DHRS3, CYP26A* and *CYP26B* in response to RA, alongside *EXOC6B* which sits adjacent to *CYP26B* on 2p13.2. Taken together, this identifies that *ATRX* IFF neuroblastoma can differentially open chromatin in response to RA at relevant gene sites.

**Figure 2.**
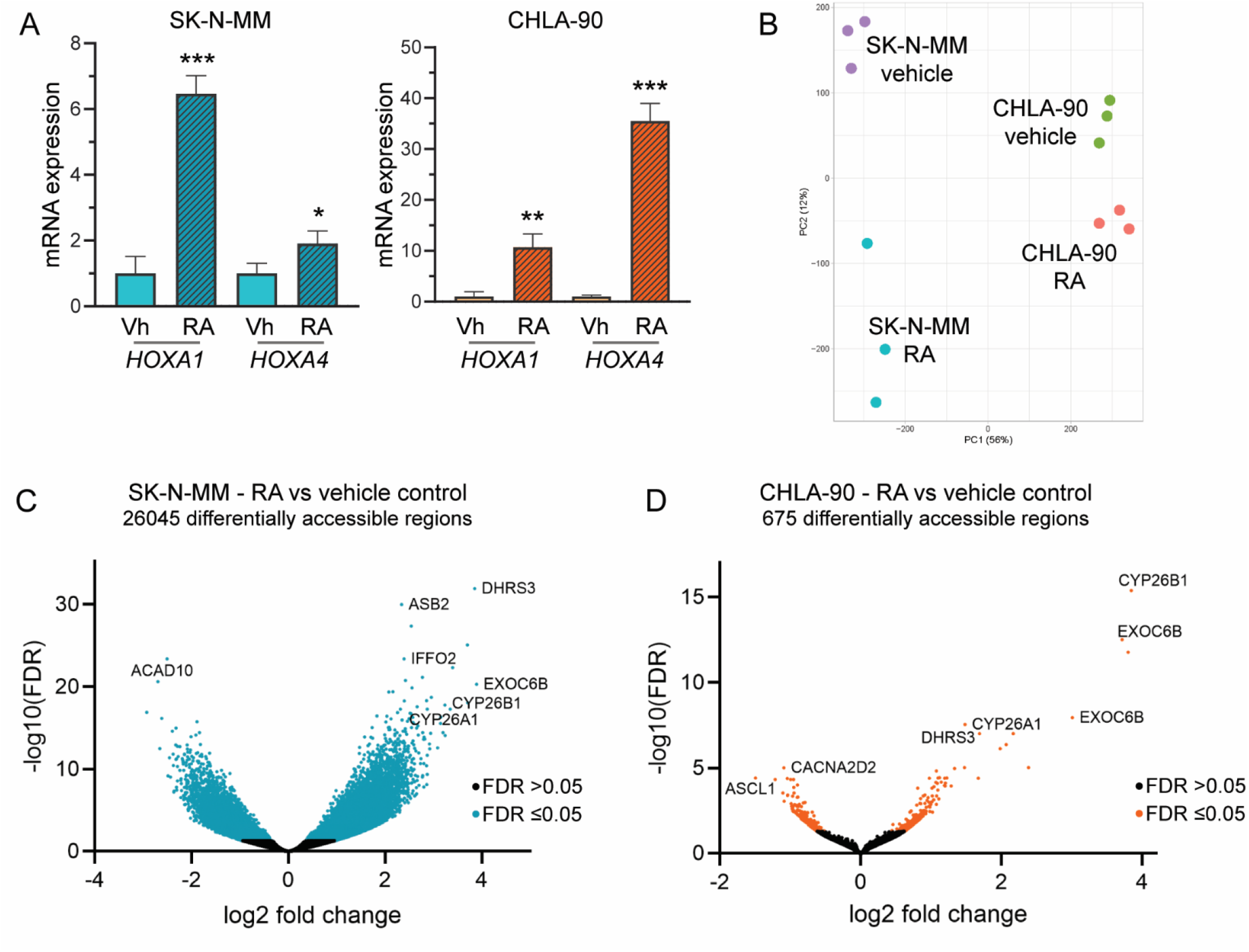
ATRX IFF neuroblastoma cell lines show increased chromatin accessibility and upregulation of direct RA target genes in response to RA. **(A)** *HOXA1* and *HOXA4* expression by RT-qPCR following RA treatment of SK-N-MM and CHLA-90 cell lines. \**P* < 0.05, \*\**P* < 0.01, \*\*\**P* < 0.001 **(B)** PCA plot showing ATAC sequencing data for 3 independent biological replicates following vehicle or RA treatment of the SK-N-MM and CHLA-90 cell lines. **(C-D)** Volcano plots showing differential accessibility in **(C)** SK-N-MM and **(D)** CHLA90 cell lines following RA treatment.

### *ATRX*-IFF neuroblastoma shows *in-vitro* sensitivity to RA

We next analysed RA sensitivity in available *ATRX* aberrant models. To our knowledge there are no currently available patient derived neuroblastoma cell lines with *ATRX* LoF. We identified a patient derived xenograft (PDX) neuroblastoma model with an *ATRX* p.Gly1748 mutation and treated this model with a clinically relevant dose of RA [18], 10 days after xenograft injection, and found no difference in tumour latency or growth in RA treated PDXs versus vehicle control (**Fig S2A**).

We next evaluated sensitivity to RA in *ATRX* IFF models. Others have shown that a non-canonical function of the ATRX-IFF in blocking neuronal differentiation results in sensitivity to EZH2 inhibition [6]. Therefore, we hypothesized that the combination of RA with EZH2 inhibition would be particularly effective for *ATRX* IFF neuroblastoma. Both ATRX IFF neuroblastoma cell lines showed sensitivity to RA as a single agent (**Fig 3A-D**) and this was further augmented with the addition of either tazemetostat or the EZH2 degrader MS1943, at doses sufficient to reduce H3K27me3 levels (**Fig 3A-E**).

**Figure 3.**
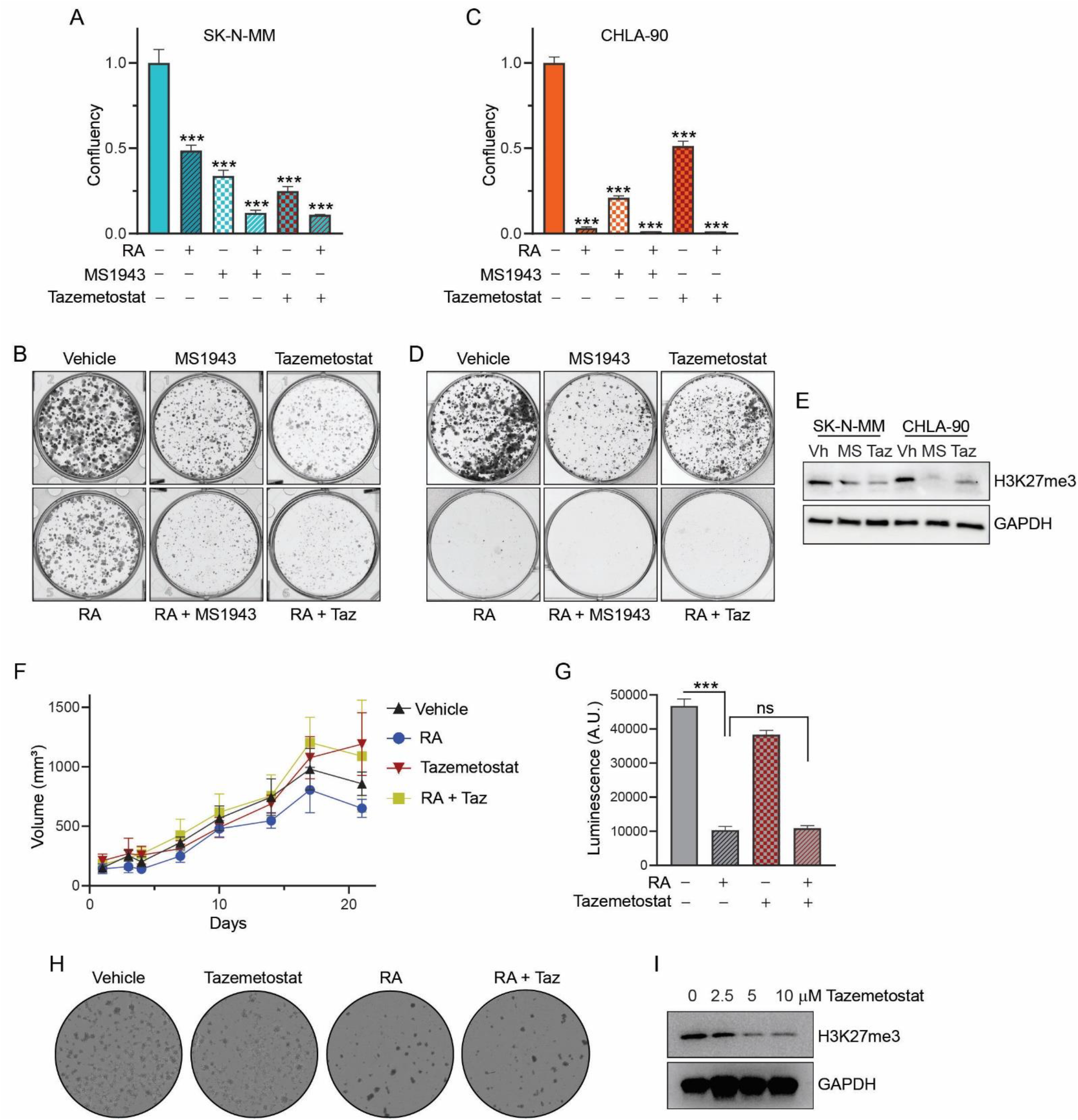
ATRX IFF neuroblastoma models show *in vitro* sensitivity to RA. Results of clonogenic assay in response to RA alone and in combination with the EZH2 inhibitors tazemetostat (taz) and MS1943 in **(A-B)** SK-N-MM and **(C-D)** CHLA-90. **(E)** Western blot for H3K27me3 levels following treatment with different EZH2i as indicated at 2.5 µM concentration. GAPDH was used as loading control. **(F)** Tumor volume of the AMC772 *in vivo* xenograft models in response to RA +/-tazemetostat. **(G-I)** *In vitro* response to RA +/-tazemetostat in the AMC772 patient derived organoid (PDO) model. **(G)** Cells were treated with 10 µM RA alone and in combination with 10 µM tazemetostat and cell viability measured via Cell Titer Glo luminescence assay. **(H)** Representative images showing cell confluency following seven days exposure to RA alone and/or in combination with 10 µM tazemetostat. **(I)** H3K27me3 levels following treatment with different doses of tazemetostat that were evaluated *in vitro*. GAPDH was used as loading control.

Given the promising *in-vitro* results in *ATRX* IFF models, we evaluated *in-vivo* sensitivity to RA alone, and in combination with tazemetostat in the AMC772 *ATRX* IFF xenograft model. However here we saw no response to either agent alone, or in combination (**Fig 3F**). Given the limitations in identifying *in-vivo* response to differentiation agents used in a minimal residual disease setting in a rapidly growing xenograft model, we assessed response to these agents in an organoid model developed from the AMC722 xenografts. Here, we identified *in-vitro* sensitivity to RA but did not see any sensitivity to tazemetostat at doses sufficient to reduce H3K27me3 levels (**Fig3 G-I**). Taken together, our results identify that *in-vitro* sensitivity to RA, but not tazemetostat is consistent across multiple *ATRX* IFF models.

## DISCUSSION

Despite increasing evidence that neuroblastoma consists of distinct molecular subgroups with different clinical phenotypes [2], all children with high-risk neuroblastoma are currently treated uniformly. RA has been used as maintenance therapy in the minimal residual disease setting for neuroblastoma for decades, with evidence of clinical benefit for some children [19]. However, it is extremely challenging to identify which patients are likely to benefit from RA given that it is used at the end of treatment in the minimal residual disease setting.

In pre-clinical studies, neuroblastoma with *MYCN* amplification has been shown to be generally sensitive to RA [20]. Here, we identify that *ATRX* IFF neuroblastoma is also generally sensitive to RA, and that ATRX IFF cell lines mount an appropriate epigenetic response to RA. In direct contrast, the epigenetic response to RA is impaired in *ATRX* LoF neuroblastoma. Our data suggests a high likelihood of RA resistance in *ATRX* LoF neuroblastoma, although the limitation of our study is that we were unable to directly evaluate sensitivity due to a lack of relevant models. Thus far, we have been unable to induce *ATRX* LoF via CRISPR-Cas9 in a RA sensitive cell line. We have also generated *ATRX* LoF clones from the NBL-S cell line [9]. However, NBL-S is highly RA resistant and does not upregulate direct target genes upon RA treatment (data not shown). This is likely to be because NBL-S carries an *NF1* mutation [21], which is has also been shown to result in RA resistance [22].

This highlights one of the significant challenges in the field, with a lack of appropriate models that reflect the heterogeneity of neuroblastoma seen in the clinic.

Here, our novel insight into the effects of *ATRX* LoF on neuroblastoma cell differentiation was triggered by an initial observation made in *ATRX* LoF iPSCs. Stem-cell models are increasingly being used to understand the underlying mechanisms of oncogenesis in neuroblastoma [23] and here, we show their potential to also give clinically relevant insight. Ongoing work in our laboratory and others to model the biological effects of specific *ATRX* alterations at later stages of neural crest differentiation will provide further insight, in addition to much needed pre-clinical models of neuroblastoma for future studies.

In conclusion, we demonstrate differences in chromatin response to RA in neuroblastoma according to type of *ATRX* alteration and propose that within the *ATRX* subgroup, there is likely to be significant heterogeneity in response to differentiation therapies depending on the precise type of alteration. This emphasises the need for improved patient stratification based on molecular features in the upfront treatment setting, alongside the need to develop novel approaches for certain subgroups of patients. Our work also highlights one of the major challenges in the field as we identify increasingly smaller molecular subgroups, each with differing underlying biology, within this already rare disease.

## METHODS

| Reagent/Resource | Reference or Source | Identifier or Catalog Number |
| --- | --- | --- |
| **Experimental Models** |  |  |
| SK0019 iPSC (wild-type and *ATRX* LoF) | Provided by Stephen Scherer, Toronto [10] | N/A |
| p53(2) cell line | George et al, EBioMedicine, 2020 [14] | N/A |
| E6 cell line | George et al, EBioMedicine, 2020 [14] | N/A |
| CHLA-90 | COG cell line repository, USA | N/A |
| SK-N-MM | Provided by Emily Bernstein, USA | N/A |
| AMC772 | Provided by Jan Molenaar, Netherlands | N/A |
| IC-pPDX17 | Gudrun Schleiermacher, France [24] | N/A |
| **Antibodies** |  |  |
| ATRX | Sigma | HPA001906 |
| GAPDH | Cell signaling | 2118 |
| H3K27me3 | Cell signaling | C36B11 |
| HRP conjugated anti-rabbit | Dako | PO448 |
| **Oligonucleotides and other sequence-based reagents** |  |  |
| HOXC4 primer | Dickson et al, 2009 [25] | N/A |
| HOXC6 primer | Dickson et al, 2009 [25] | N/A |
| HOXC9 primer | Dickson et al, 2009 [25] | N/A |
| ATAC primers | Corces et al, 2017 [26] | N/A |
| **Chemicals, Enzymes and other reagents** |  |  |
| NANOG taqman probe | ThermoFisher Scientific | Hs02387400_g1 |
| OCT4 taqman probe | ThermoFisher Scientific | Hs04260367_gH |
| SOX2 taqman probe | ThermoFisher Scientific | Hs04234836_s1 |
| CDX2 taqman probe | ThermoFisher Scientific | Hs01078080_m1 |
| CYP26A taqman probe | ThermoFisher Scientific | Hs00175627_m1 |
| HOXA1 taqman probe | ThermoFisher Scientific | Hs05047045_s1 |
| HOXA4 taqman probe | ThermoFisher Scientific | Hs01573270_m1 |
| GAPDH taqman probe | ThermoFisher Scientific | Hs02786624_g1 |
| PowerUp™ SYBR™ Green master mix for qPCR | ThermoFisher Scientific | A25743 |
| Taqman™ Universal PCR master mix | ThermoFisher Scientific | 4304437 |
| Recombinant Human FGF basic (146 aa) protein CF | R&D systems | #233-FB/CF |
| CHIR99021 | Tocris | \|4423 |
| Geltrex basement membrane matrix | ThermoFischer | A1413201 |
| Tagment DNA enzyme and buffer kit | Illumina | 20034197 |
| 13-cis-retinoic acid | Sigma Aldrich | R3255 |
| Tazemetostat | Selleckchem | S7128 |
| MS1943 | MedChemExpress | HY-133129 |
| Maxima First Strand cDNA synthesis kit for RT-qPCR | ThermoFisher Scientific | K1641 |
| ACK lysis buffer | Thermofischer | A1049201 |
| GlutaMAX™ | Thermofischer | 21885025 |
| Ham's F-12 Nutrient Mix | Thermofischer | 15344967 |
| B-27 Supplement™ (50X) | Thermofischer | 11530538 |
| minus vitamin A |  |  |
| N-2 supplement | ThermoFischer | 17502048 |
| Recombinant Human EGF | PeproTech | AF-100-15 |
| recombinant Human FGF-basic | PeproTech | AF-100-18B |
| recombinant Human IGF-I | PeproTech | 100-11-100UG |
| recombinant Human PDGF-AA and PDGF-BB | PeproTech | #100-13A<br>#100-14B |

### Cell culture and differentiation

Paired *ATRX* wild-type and LoF SK0019 iPSC’s were generated in the Scherer laboratory [10]. iPSCs were cultured on Geltrex™ reduced growth factor basement membrane and maintained in StemFlex™ media and differentiated into human axial progenitor cells using published protocols [11]. The neuroblastoma cell lines p53(2), E6, CHLA-90 and SK-N-MM were maintained in 10% fetal calf serum Dulbecco’s Modified Eagle Medium (DMEM). The AMC772 organoid cell line was generated from xenografts established in the adrenal gland of NSG mice. Tissue was minced and strained using 70 μm cell strainer. Cells were then washed with ACK lysis buffer for 5 minutes to remove red blood cells before inactivation using Low-glucose DMEM + 10% fetal calf serum. Cells were again washed in 10 mL DMEM low glucose GlutaMAX™. Cells were maintained in organoid media (DMEM low glucose GlutaMAX™ Filemented with: 25% Ham’s F-12 Nutrient Mix, B-27™ Supplement, N-2 Supplement, 100 U/mL penicillin and 100 μg/mL streptomycin, 40 ng/mL recombinant human EGF, 200 ng/mL recombinant human FGF-basic, 10 ng/mL recombinant human IGF-I, 10 ng/mL recombinant human PDGF-AA and PDGF-BB. 13-cis-retinoic acid, tazemetostat and MS1943 were all dissolved in dimethyl-sulfoxide (DMSO) to make 10 mM stock concentrations.

### Quantification of mRNA by RT-qPCR

RNA was isolated from cells using the Qiagen RNeasy kit and cDNA synthesized using the Maxima First Strand cDNA synthesis kit (ThermoFisher). Either commercially available Taqman probes or SYBR green probes with published sequences were used as indicated in the reagents table. qPCR reactions were performed as per manufacturer’s instructions using the QuantStudio3™ real time PCR machine. *GAPDH* was used as a housekeeping gene. Relative mRNA abundance of target genes was calculated by the 2(−ΔΔCq) method. Relative quantity compared with control was calculated and is displayed except for *HOXA1* and *HOXA4* expression in p53(2) and E6 cell lines where baseline levels were not detectable in untreated cells for comparison hence inverse log of delta CT is displayed. All RT-qPCR data is representative of at least 2 independent experiments, each performed in triplicate.

### ATAC sequencing

Neuroblastoma cell lines were treated with either vehicle control (DMSO) or 10 µM RA for 72 hours, before collection for ATAC sequencing. 25 000 cells were used per experiment and ATAC sequencing libraries were prepared using the OMNI-ATAC protocol [26], with modifications as previously published [27]. All ATAC-sequencing experiments were performed in biological triplicates. ATAC-sequencing was carried out at the Genomics facility at the Francis Crick Institute.

### ATAC sequencing data analysis

All ATAC sequencing data analyses were performed on the Crick HPC cluster using Nextflow v22.04.0 [28] and Singularity v3.6.4 [29]. The nf-core/atacseq pipeline (release 3.0) [28] was executed with the Crick profile for alignment, peak calling and quality control. The GRCh38, Ensembl release 95 genome reference was used. Consensus peaksets were imported into R [30] and analysed with DiffBind [31]. Counts were generated then normalised for library size and background. A generalised linear model incorporating Tissue, Treatment, and their interaction was specified. Normalised read counts for peaks were log_2_-transformed and principle component analysis (PCA) was performed on the transposed matrix using prcomp [30]. The percentage variance explained by PC1 and PC2 was calculated from the standard deviations. Scores were plotted with ggplot2 [32].

All peaks were annotated to the nearest genomic feature using ChIPpeakAnno [33]. Distance to TSS was calculated from peak midpoint. Peaks were annotated to nearest promoters using the Ensembl GRCh38.95 GTF. Promoter regions were defined as ±5 kb around the TSS. Overlaps between peak ranges and promoter windows were detected using IRanges::findOverlaps [34].

Differential binding results were retrieved, and a data frame was constructed with log_2_ fold-change (x-axis), –log_10_FDR (y-axis), and a significance flag (FDR ≤ 0.05). Points exceeding |log_2_ fold-change| > 1 and –log_10_FDR > 1 were labelled with gene symbols using ggrepel [35]. Volcano plots were drawn with ggplot2 [32].

For gene ontology (GO) enrichment, for each comparison, DiffBind output (Fold and FDR) was parsed into separate “up” and “down” gene lists. GO Biological Process enrichment was computed with clusterProfiler [36]. From the list of GO biological processes, redundant terms were eliminated using REVIGO (http://revigo.irb.hr/) [17]. A small list of GO terms was obtained and graphs were drawn with ggplot2.

Motif enrichment analysis was performed using position weight matrices (PWMs) for vertebrate transcription factors which were retrieved from JASPAR2020 [37]. Peaks were scanned for motif enrichment using monaLisa [38]: sequences were extracted from the GRCh38 genome fasta using Biostrings [39], binned by log2 fold-change into three bins (–10 to 0, 0–10), and motif enrichment calculated with calcBinnedMotifEnrR from the monaLisa package. Motifs with |log2 enrichment| > fold-change threshold and –log10 padj > threshold were retained. Similar motifs were clustered by Pearson similarity of PFMs and visualized as heatmaps with sequence-logos.

### Clonogenic assays

For adherent cell lines 1000-10000 cells were plated in 2 ml medium, in triplicate in 6-well plates. After 24 hours medium was exchanged and drugs or DMSO added and cells cultured for 1–3 weeks, with fresh drug containing media being replaced twice weekly. 10 µM RA was used for all cell lines, except for CHLA-90 which was treated with 100 nM RA for the clonogenic assay, CHLA-90 and SK-N-MM cell lines were treated with 5 µM of either tazemetostat or MS1943. Adherent cells were stained by 0.01% crystal violet in distilled water for 30 minutes at room temperature, then washed before drying of the plates overnight. Plates were visualized using the gelcount machine (Oxford Optronix). Confluency was calculated using imageJ processing software.

Cell Titer-Glo 3D was used to assess treatment response in the AMC772 organoid model. A total of 20000 cells were plated in triplicate in 100 μL of organoid media in an opaque 96-well plate and incubated for 4 days in humidified incubator at 37°C with 5% CO_2_ to allow spheroid formation. Following spheroid formation, the media was replaced with 100 μL of fresh media containing either DMSO, Tazmetostat and/or RA, then incubated for an additional seven days, with imaging performed on the Opera Phenix™ on days 1, 3, 5, and 7 post-treatments. On day 7, 100 μL CellTiter-Glo® 3D Reagent (Promega Cat#G9681) was added to each well mixed on an orbital shaker for 10 minutes then allowed to stabilize for 30 minutes before fluorescence was measured at 560 nm using a spectrometer.

### Western blot

SK-N-MM and CHLA-90 were treated with tazemetostat and MS-1943 at a concentration of 2.5 µM for 72 hours then proteins were isolated by Pierce IP Lysis buffer containing phosphatase inhibitor cocktail.

For the AMC772 organoids, 200000 cells were seeded in triplicate into 6-well plates with 2.5 mL of organoid media and incubated for 4 days in humidified incubator at 37°C with 5% CO_2_ to allow spheroid formation. Following spheroid formation, the media was replaced with 2.5 mL of fresh media containing either DMSO, tazmetostat and/or RA. Cells were then incubated for an additional seven days before cells lysis in 1X RIPA lysis buffer supplemented with a protease inhibitor and phosphatase inhibitor cocktail.

Protein concentrations were measured by BCA protein assay. Membranes were incubated overnight with primary antibodies, washed with TBS-T and then incubated 1 hour at room temperature with the secondary antibody listed in the reagents table. Membranes were incubated with Clarity™ Western ECL Substrate before imaging using the ChemiDoc™ Imaging System (Bio-Rad).

### *In-vivo* experiments

AMC772 xenografts were established subcutaneously in NOD scid gamma (NSG) mice as published previously [40]. Tumour development was monitored by palpation by an experienced technician and animals randomly allocated to treatment arms once a tumour was palpable. The clinically relevant dose of 53 mg/kg 13-*cis*-retinoic acid was used [18], administered via oral gavage once daily, Monday to Friday, dissolved in a total volume of 200 µL corn oil. Tazemetostat was administered at a dose of 10 mg/kg intraperitoneally once daily, Monday–Friday, in 50 µL total volume of 5% DMSO (+4.5% PEG300, 5% tween and ddH20). Mice were housed in autoclaved, aseptic cages in specific pathogen-free rooms in The Institute of Cancer Research (ICR) animal facility and allowed access to sterile food and water ad libitum. Experiments were approved by The ICR Animal Welfare and Ethical Review Body and performed in accordance with the UK Home Office Animals (Scientific Procedures) Act 1986, the UK National Cancer Research Institute guidelines for the welfare of animals in cancer research and the ARRIVE (animal research: reporting *in-vivo* experiments) guidelines.

The IC-pPDX17 was established from a NB tumour biopsy following informed consent from the parents, as published previously [24, 41].Swiss Nude mice were engrafted with tumours in their interscapular fat pad on day 0 (D0). Treatment with 13cRA or vehicle injection began on day 10 (D10) after tumour engraftment. The mice then received daily intraperitoneal injections of either RA or vehicle control 5 days per week for three weeks. Mouse weight and tumour size were monitored three times per weekly. Mice were euthanized when tumours reached a predetermined ethical size limit of approximately 1,500-2,000 mm^3^. All procedures were approved by the Institutional Review Board of the Institut Curie. All animal experiments complied with current European/French legislation (articles R.214-87 to R.214-126 of the Decree n°2013-118 of February 1st) and were carried out in accredited animal facilities of the Institut Curie.

## Acknowledgements

We would like to thank Stephen Scherer for sharing the *ATRX* WT and LoF IPSC’s generated in his laboratory and Tom Frith for his help and support with IPSC neural crest differentiation protocols.

FL and SLG are supported by a Cancer Research UK Clinician Scientist Fellowship. MS, LC and SLG are supported by ICR HEFCE funding. LD is supported by a CCLG Little Princess Trust Project Grant. VT is supported by funding from the ArcoBaleno Trust/Siobhan’s Superstar legacy. KB is supported by the nMGN Cancer Cluster, MRC KG and BC are supported by Chesler CR UK Prog Award A28278. KC is also supported by the Brain Tumour Charity (GN-000690).

In France, this work was supported by the Annenberg Foundation, the Association Hubert Gouin Enfance et Cancer, Imagine for Margo, the Fondation ARC pour la Recherche contre le Cancer (ARC), and Fights Kids Cancer. Establishment of PDX models was performed under the MAPPYACTS protocol (clinicaltrial.gov: NCT02613962). Molecular characterization of the PDX models was supported within the framework of the IMI2 ITCC P4 program funded by IMI2 grant agreement no. 116064 ITCC-P4.

## SUPPLEMENTARY FIGURES

**Supplementary Fig 1:**
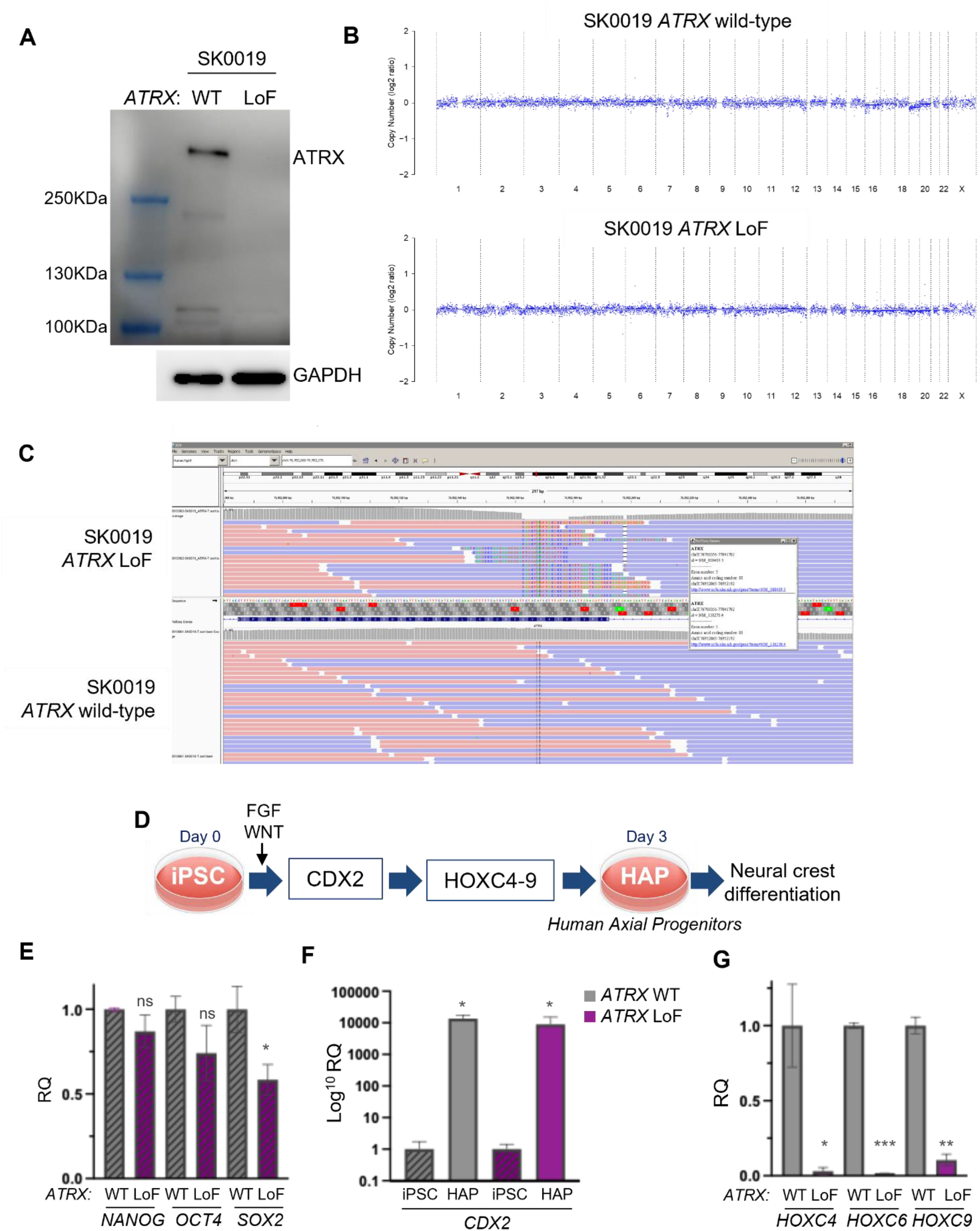
**(A)** ATRX protein expression in paired SK0019 *ATRX* wild-type and LoF iPSC lines. **(B)** Confirmation of normal karyotype in SK0019 WT and *ATRX* LoF iPSCs by low coverage whole genome sequencing and **(C)** detection of stop codon in exon 5 of *ATRX* by NGS panel sequencing. **(D)** Experimental scheme for differentiation of iPSCs into human axial progenitor cells (HAPs). **(E)** *NANOG, OCT4* and *SOX2* mRNA expression by RT-qPCR relative to *GAPDH* in SK0019 iPSCs, comparing relative quantity in *ATRX* LoF SK0019 iPSC to matched WT SK0019 iPSCs (*t* test, \**P* < 0.05). **(F)** *CDX2* expression by RT-qPCR relative to *GAPDH*, in SK0019 HAPs relative to SK0019 iPSCs (*t* test, \**P* < 0.05). **(G)** *HOXC4, HOXC8* and *HOXC9* expression relative to *GAPDH*, comparison of WT HAPs with *ATRX* LoF HAP (*t* test, \**P* < 0.05, \*\**P* < 0.01, \*\*\**P* < 0.001).

**Supplementary Fig 2:**
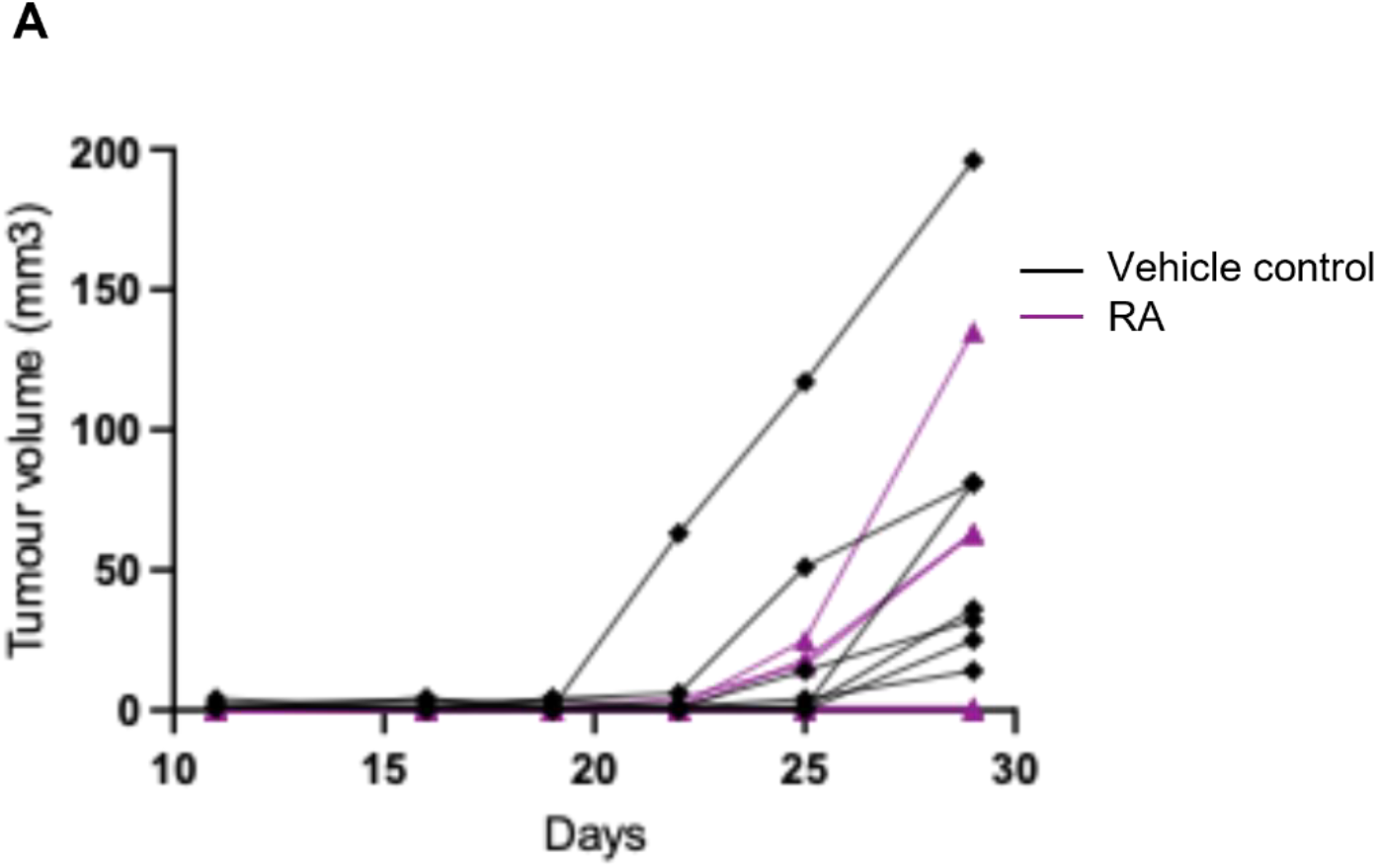
**(A)** The IC-pPDX17 PDX (containing an *ATRX* p.Gly1748Arg mutation) was engrafted at day 0 and either RA 53mg/kg or vehicle treatment commenced at D11 post engraftment and tumour volume measured.

## References

1. Ackermann, S., et al., A mechanistic classification of clinical phenotypes in neuroblastoma. Science, 2018. 362(6419): p. 1165–1170.

2. Zeineldin, M., et al., MYCN amplification and ATRX mutations are incompatible in neuroblastoma. Nat Commun, 2020. 11(1): p. 913.

3. Cheung, N.K., et al., Association of age at diagnosis and genetic mutations in patients with neuroblastoma. JAMA, 2012. 307(10): p. 1062–71.

4. Aguilera, P. and A.J. Lopez-Contreras, ATRX, a guardian of chromatin. Trends Genet, 2023. 39(6): p. 505–519.

5. van Gerven, M.R., et al., Mutational spectrum of ATRX aberrations in neuroblastoma and associated patient and tumor characteristics. Cancer Sci, 2022. 113(6): p. 2167–2178.

6. Qadeer, Z.A., et al., ATRX In-Frame Fusion Neuroblastoma Is Sensitive to EZH2 Inhibition via Modulation of Neuronal Gene Signatures. Cancer Cell, 2019. 36(5): p. 512–527 e9.

7. van Gerven, M.R., et al., Two opposing gene expression patterns within ATRX aberrant neuroblastoma. PLOS ONE, 2023. 18(8): p. e0289084.

8. Pugh, T.J., et al., The genetic landscape of high-risk neuroblastoma. Nat Genet, 2013. 45(3): p. 279–84.

9. Lorenzi, F., et al., ATRX mutations mediate an immunogenic phenotype and macrophage infiltration in neuroblastoma. Cancer Lett, 2025: p. 217495.

10. Deneault, E., et al., Complete Disruption of Autism-Susceptibility Genes by Gene Editing Predominantly Reduces Functional Connectivity of Isogenic Human Neurons. Stem Cell Reports, 2018. 11(5): p. 1211–1225.

11. Frith, T.J.R. and A. Tsakiridis, Efficient Generation of Trunk Neural Crest and Sympathetic Neurons from Human Pluripotent Stem Cells Via a Neuromesodermal Axial Progenitor Intermediate. Curr Protoc Stem Cell Biol, 2019. 49(1): p. e81.

12. Young, T., et al., Cdx and Hox genes differentially regulate posterior axial growth in mammalian embryos. Dev Cell, 2009. 17(4): p. 516–26.

13. Lengerke, C., et al., The cdx-hox pathway in hematopoietic stem cell formation from embryonic stem cells. Ann N Y Acad Sci, 2007. 1106: p. 197–208.

14. George, S.L., et al., Therapeutic vulnerabilities in the DNA damage response for the treatment of ATRX mutant neuroblastoma. EBioMedicine, 2020. 59: p. 102971.

15. Balmer, J.E. and R. Blomhoff, Gene expression regulation by retinoic acid. J Lipid Res, 2002. 43(11): p. 1773–808.

16. Gomez, R.L., et al., Super-enhancer associated core regulatory circuits mediate susceptibility to retinoic acid in neuroblastoma cells. Front Cell Dev Biol, 2022. 10: p. 943924.

17. Supek, F., et al., REVIGO summarizes and visualizes long lists of gene ontology terms. PLoS One, 2011. 6(7): p. e21800.

18. Nolting, S., et al., Combination of 13-Cis retinoic acid and lovastatin: marked antitumor potential in vivo in a pheochromocytoma allograft model in female athymic nude mice. Endocrinology, 2014. 155(7): p. 2377–90.

19. Matthay, K.K., et al., Treatment of high-risk neuroblastoma with intensive chemotherapy, radiotherapy, autologous bone marrow transplantation, and 13-cis-retinoic acid. Children’s Cancer Group. N Engl J Med, 1999. 341(16): p. 1165–73.

20. Zimmerman, M.W., et al., Retinoic acid rewires the adrenergic core regulatory circuitry of childhood neuroblastoma. Sci Adv, 2021. 7(43): p. eabe0834.

21. Izquierdo, E., et al., Development of a targeted sequencing approach to identify prognostic, predictive and diagnostic markers in paediatric solid tumours. Oncotarget, 2017. 8(67): p. 112036–112050.

22. Holzel, M., et al., NF1 is a tumor suppressor in neuroblastoma that determines retinoic acid response and disease outcome. Cell, 2010. 142(2): p. 218–29.

23. Saldana-Guerrero, I.M., et al., A human neural crest model reveals the developmental impact of neuroblastoma-associated chromosomal aberrations. Nat Commun, 2024. 15(1): p. 3745.

24. Boeva, V., et al., Heterogeneity of neuroblastoma cell identity defined by transcriptional circuitries. Nat Genet, 2017. 49(9): p. 1408–1413.

25. Dickson, G.J., T.R. Lappin, and A. Thompson, Complete array of HOX gene expression by RQ-PCR. Methods Mol Biol, 2009. 538: p. 369–93.

26. Corces, M.R., et al., An improved ATAC-seq protocol reduces background and enables interrogation of frozen tissues. Nat Methods, 2017. 14(10): p. 959–962.

27. Lattke, M., et al., Extensive transcriptional and chromatin changes underlie astrocyte maturation in vivo and in culture. Nat Commun, 2021. 12(1): p. 4335.

28. Ewels, P.A., et al., The nf-core framework for community-curated bioinformatics pipelines. Nat Biotechnol, 2020. 38(3): p. 276–278.

29. Kurtzer, G.M., V. Sochat, and M.W. Bauer, Singularity: Scientific containers for mobility of compute. PLoS One, 2017. 12(5): p. e0177459.

30. R: A language and environment for statistical computing. R Foundation for Statistical Computing. https://www.R-project.org/ 2023.

31. Stark, A.B. G.D., DiffBind: differential binding analysis of ChIP-seq peak data. Bioinformatics, 2011. 27.

32. Wickham, H., ggplot2: Elegant Graphics for Data Analysis. Springer-Verlag New York, 2016.

33. Zhu, L.J., et al., ChIPpeakAnno: a Bioconductor package to annotate ChIP-seq and ChIP-chip data. BMC Bioinformatics, 2010. 11: p. 237.

34. Lawrence, M., et al., Software for computing and annotating genomic ranges. PLoS Comput Biol, 2013. 9(8): p. e1003118.

35. Slowikowsi, K., ggrepel: Automatically position non-overlapping text labels with ‘ggplot2’. R package version 0.9.1. 2021.

36. Yu, G., et al., clusterProfiler: an R package for comparing biological themes among gene clusters. OMICS, 2012. 16(5): p. 284–7.

37. Fornes, O., et al., JASPAR 2020: update of the open-access database of transcription factor binding profiles. Nucleic Acids Res, 2020. 48(D1): p. D87–D92.

38. Krebs, A.R.B. M.; Starlinger, J.; Lähnemann, D.; □□□□□, W., monaLisa: a fast and flexible tool for motif analysis in large-scale sequencing data. Bioinformatics, 2021. 37.

39. Pages, H.A. P.; Gentleman, R.; & DebRoy, S., Biostrings: String objects representing biological sequences, and matching algorithms. R package version 2.68.1., in R package version 2.68.1. 2023.

40. George, S.L., et al., A tailored molecular profiling programme for children with cancer to identify clinically actionable genetic alterations. Eur J Cancer, 2019. 121: p. 224–235.

41. Marques Da Costa, M.E., et al., A biobank of pediatric patient-derived-xenograft models in cancer precision medicine trial MAPPYACTS for relapsed and refractory tumors. Commun Biol, 2023. 6(1): p. 949.

